# Atrazine induced *In vivo* immunotoxicity in Bivalves

**DOI:** 10.1101/2021.06.29.450311

**Authors:** A.N Muhammed Zafar Iqbal, M. Azhar Iqbal Navalgund

## Abstract

Atrazine is ubiquitously used broad-spectrum herbicide to control the weeds in agriculture. The present study aimed to evaluate the acute toxicity and immunotoxicity of Atrazine in two ecologically and economically important bivalves. Acute toxicity of atrazine evaluated in triplicates by taking control and six experimental groups each comprising of 30 animals and treated with a range of atrazine from 2 PPM to 12 PPM for 96 hours. Mortalities were recorded for every 24 hours until 96 hours and data analyzed by one-way ANOVA and Dunnett T-test. The results indicated a significant increase in mortalities with increase in dose and time of exposure in both species. The values of LC_50_ were determined as 6.10 PPM and 4.90 PPM respectively for *Perna viridis* and *Paphia malabarica*. Furthermore, the immunotoxic potential of atrazine assessed by treating mussels and clams with the five sub-lethal doses of atrazine for 14 days and quantifying the viability of hemocytes by using simple yet reliable Tryphan blue exclusion assay. The results of the present study suggest atrazine-induced immunotoxicity by decreasing the number of viable hemocytes in bivalves. Hemocytes with phagocytic function are indispensable to confer innate immunity in bivalves, decreased viability of these cells leads to compromised immunity. This study is first of its kind to implicate atrazine with the immunotoxicity in bivalves.

## Introduction

Agriculture produce has enormously increased by employing several strategies; one such strategy is to get rid of weeds by using herbicides, to help the crops get maximum benefit from available resources. Different classes of herbicides use different mechanism of actions, atrazine is a broad-spectrum herbicide, which interferes with the proteins of PS-II of photosynthesis to control the weeds (Trebst, 2008). Ubiquitous use of atrazine is going on since from more than 60 years in paddy, sugar cane, sorghum and other horticulture crops (Müller, 2008). Its efficacy and cost encourages the farmers to use it regularly in large quantities. Atrazine is most effective if used in recommended doses but unfortunately, excess application is paving way for natural selection and emergence of resistant weed varieties (Müller, 2008). International survey of herbicide resistance reported development of resistance in 69 weed species, second only to ALS inhibitors (Heap, 2014). Atrazine is most commonly used herbicide throughout the world including India and US (Short and Colborn, 1999). Large-scale use of atrazine makes it one of the most common contaminant of surface waters and can persist in environment for a considerable period of time (Koskinen and Clay, 1997). Atrazine already reported to be a substance with reproductive and immunotoxic risk for number of animal models (Cummings et al., 2000; Stoker et. al. 2002; Laws et.al. 2003; Whale et al., 2003; Filipov et al., 2005; Opute et. al. 2021). Sustainable use of this herbicide needs proper evaluation from other animal models as well as various other possible kinds of risk perspectives associated with it. Toxic effects on non-target organisms is really of great interest and relevance, especially with aquatic animals.

Ubiquitous use of atrazine has the potential to bring about the undesirable and precarious effects on several ecosystems, aquatic ecosystems being the most vulnerable amongst all (Flynn & Spellman 2009). Agricultural chemicals contaminate surface water locally but have the potential to impact at global scale by entering into the food chain or by bioaccumulation (Shomar et. al. 2006). Acute and chronic toxicities of the pesticides and herbicides to the non-target aquatic organisms is a well-known fact through the scientific investigations (LeBlanc et al. 1997; María Elena et al. 2007; Palma et. al. 2008; Flynn & Spellman 2009; Ralston-Hooper et. al. 2009; Nwani et al. 2010; Tongo and Ezemonye, 2015; Ullah et al. 2015; Klementová et. al. 2019; Westlund et al. 2018). The ecological risk potentials associated with atrazine on the aquatic weeds and fishes is studied extensively (Solomon et al. 1996; Giddings et al., 2005; Glen et al., 2014). Most of the investigators have focused their attention on aspects such as metabolism, oxidative stress, lipid peroxidase activity, antioxidant enzyme systems (Ullah et. al. 2015; Nwani et. al. 2010; Ralston-Hooper et. al. 2009). Some studies have also investigated the role of atrazine in aggregation behavior of fresh water mussels (Flynn and Spellman, 2009). As the aquatic animals are more vulnerable to the toxic effects of atrazine, due attention is required to investigate the acute and chronic toxicity. Bivalves are one of the most susceptible group of animals, which experience maximum loads of environmental contaminants due to their intertidal and estuarine habitats. Moreover, bivalves are also known to accumulate heavy metals and pollutants in their soft tissues and some even have the ability to transform xenobiotics into Mutagens (Sezer et al. 2020)

Bivalves such as mussels, oysters, clams are an important part of human diet with annual consumption of more than 90000 t per annum. However, the risk assessment of contaminants in these groups of animals is not as it should have been. There are few reports on the toxicity of heavy metals and genotoxic effects of some environmental mutagenic agents. (Prakash and Jagannath Rao, 1995; Sokolova, 2004). Several other areas of toxicological risk assessment is yet to be investigated and reported.

Invertebrates rely solely on the innate immunity and hemocytes are cells, which play a critical role in internal defense (Canesi and Pruzzo, 2016; Burgos-Aceves and Faggio, 2017; Destoumieux-Garzón et al. 2020). Molluscan hemocytes have been described in detail and their classification based on their functions (Cheng, 1984). Besides their primary role as cells of immunity, these cells are also known to perform several other functions (Cheng, 1977). Hence, the hemocytes are indeed most important cells and deserve greater scientific attention for their comprehensive understanding

In the present study, we are assessing the toxic effects of atrazine on the hemocytes of two edible bivalves viz; green mussel, *Perna viridis* (Order Mytilida) and clam, *Paphia malabarica* (Order Venerida) by Tryphan Blue Exclusion Assay (Strober, 2014). This technique is very sensitive and reliable for quantification of the viable cells under any simple laboratory set up. Distinction between viable and non-viable cells can be done by observing clear / blue cytoplasm under light microscope. Other assays such as MTT Assay, MTS assay, Annexin V assay, TUNEL assay or caspase assays are more sensitive but expensive, generally used in apoptosis and cancer research. Hence, the Tryphan blue exclusion assay is most suitable and precise method for studying viability of the suspended hemocytes.

## Materials and Method

### *Perna viridis* (Order: Mytilidae)

#### Animal Collection and acclimatization

Collected fully-grown and mature *Perna viridis* from designated field stations during low tide and immediately transported mussels to the laboratory. Mussels were acclimatized under laboratory conditions with proper aeriation set up for five days by following suitable aquaculture practices (Muhammed Zafar Iqbal et al. 2008). During this time, animals fed ad libitum on the phytoplankton, *Chaetocerous sp*.

#### 96 hour LC_50_ Calculations and sub-lethal doses

For the determination of the median lethal concentration (LC_50_) carried out by 96 hour experiment, 210 healthy animals divided into six experimental groups with 30 mussels each and one control group with 30 mussels, treated with 0 (Control), 2, 4, 6, 8, 10 and 12 PPM of atrazine (TATA-Atrataf 50% WP) for a period of 96 hours. Mortality was recorded at 24, 48, 72 and 96 hours and data tabulated for further analysis. The experiments of acute toxicity conducted in triplicates for better accuracy. Data analyzed statistically by one way ANOVA to find out the significant difference of mortality between control and experimental groups. When the difference between the groups found to be significant further analysis was done by Dunnett T test to compare control group with each of the experimental groups. Based on the results, lowest observed effective level (LOEL) was noted. Acute toxicity data was also analyzed to determine the median lethal concentration (LC_50_) and 95 % confidence Interval by regression analysis. All statistical analysis were done by using IBM SPSS 24.0.

Immunotoxic effects of atrazine was studied by exposing the green mussels to five sub-lethal doses taken between 1/25^th^ to 1/5^th^ values of median lethal concentrations (LC_50s_). Each dose was prepared by dissolving the required amount of atrazine in 150 ml tap water by stirring continuously at low speed on a magnetic stirrer for 2 hours.

#### Exposure of green mussels to sub-lethal doses

Fifty healthy and fully-grown green mussels from acclimatized stock introduced in each of the six large fiber containers with 100 liters of brackish water. Freshly prepared doses of 0.25 (1/25th of LC_50_), 0.50 (1/12th), 0.75 (1/7th), 1.00 (1/6th) and 1.25 (1/5^th^) decanted one each into the five experimental group containers. Concurrently, control group was maintained without any atrazine

#### Hemolymph Extraction

Experiment was carried out for 14 days and the hemolymph was extracted on 2, 4, 6, 8, 10, 12 and 14 days from posterior adductor muscles of 5 to 6 green mussels each from control, 0.25, 0.50, 0.75, 1.00 and 1.25 PPM by using 5 cc syringe.

#### Tryphan Blue Exclusion Assay

Tryphan Blue Exclusion Assay was performed by following the protocol provided by Sigma – Aldrich. Cell suspension of 10 ml was prepared by taking 2 ml of hemolymph, 3 ml of Hank’s Balanced Salt Solution (HBSS) and 5 ml of Tryphan Blue w/v (Sigma-Aldrich – T8154). The cell suspension was allowed to stand for 5 to 7 minutes for proper staining. Cover glass was placed on the clean Neubauer’s Hemocytometer and small quantity of cell suspension was allowed with the help of pipette to flow onto the chamber of the hemocytometer by capillary action. Cells in four corner 1 mm^2^ and one central 1 mm^2^ observed, viable cells with clear cytoplasm and nonviable cells with blue coloured cytoplasm recorded separately. The procedure was repeated for the second chamber as well. More than 2000 Viable and nonviable cell counts were recorded from each of the six experimental (0.25, 0.50, 0.75, 1.00 and 1.25 PPM) and control groups by following the same procedure on every alternate day (2, 4, 6, 8, 10, 12 and 14^th^ days).

#### Calculations of viable and nonviable hemocytes

Hemocytes per ml, number of hemocytes, percent viability and percent nonviability were calculated by using following formulas (As per protocol given by Sigma-Aldrich)

1. Hemocytes per ml = Average count per square × 5 × 10^4^
2. Total hemocytes = Hemocytes / ml × 10 ml
3. Viable Hemocytes Count (VHC) in % = Unstained (Viable) / Total cells ×100
4. Non-Viable Hemocytes Count (NVHC) in % = Stained cells (Nonviable)/Total cells ×100

### Statistical Analysis

Data of percent viability and percent non-viability analyzed statistically to determine the mean lethal concentration (LC_50_) value with 95 % confidence Interval by applying Regression test by using IBM SPSS 24.0.

### *Paphia malabarica* (Order Veneroida)

#### Animal Collection and acclimatization

Clams, *Paphia malabarica* were collected from sediments at Kali estuary (14°50□39.60□N 74°07□31.99□E) and immediately transported to laboratory. Acclimatized and fed the clams as described for *Perna viridis*.

#### 96 hour LC_50_ Calculations and sub-lethal doses

Procedure as described for green mussels

#### Exposure of clams to sub-lethal doses

As the yield of hemolymph is less, to get sufficient quantity, one hundred clams were introduced in each of six large fiber containers with 100 liters of brackish water. Freshly prepared doses of 0.20 PPM, 0.40 PPM, 0.60 PPM, 0.80 PPM and 1.00 PPM decanted one each into the five containers. Concurrently, control maintained without any atrazine under same conditions.

#### Hemolymph Extraction

As described for green mussels

#### Tryphan Blue Exclusion Assay

As described for green mussels

#### Calculations of viable and nonviable cells

As described for green mussels

#### Statistical analysis

As described for green mussels

## Results

### Perna viridis

#### Acute toxicity studies

Acute toxic effects of atrazine never been studied on the green mussels (*Perna viridis*) and clams (*Paphia malabarica*). We are reporting the median lethal concentration (LC_50_) of atrazine for these two species for the first time. Median lethal concentration (LC_50_) and lowest observed effective level (LOAL) for were evaluated by conducting 96 hour experiments in triplicates. Mortality was recorded at 24, 48, 72 and 96 hours for the experimental and control groups.

The results of the experiments analysed by one way ANOVA indicated significant difference in mortality between control and experimental groups with F = 79.64 and P < 0.0001, further analysis of multiple comparisons by Dunnett T test between control group with each of the experimental groups also found to be significant with P < 0.0001. The lowest observed effective level (LOEL) was found to be 2.00 PPM (P < 0.009) with 95 % confidence interval (1.18, 8.81). The results of one way ANOVA and Dunnett T test are presented in **Table 1**.

**Table 1.**
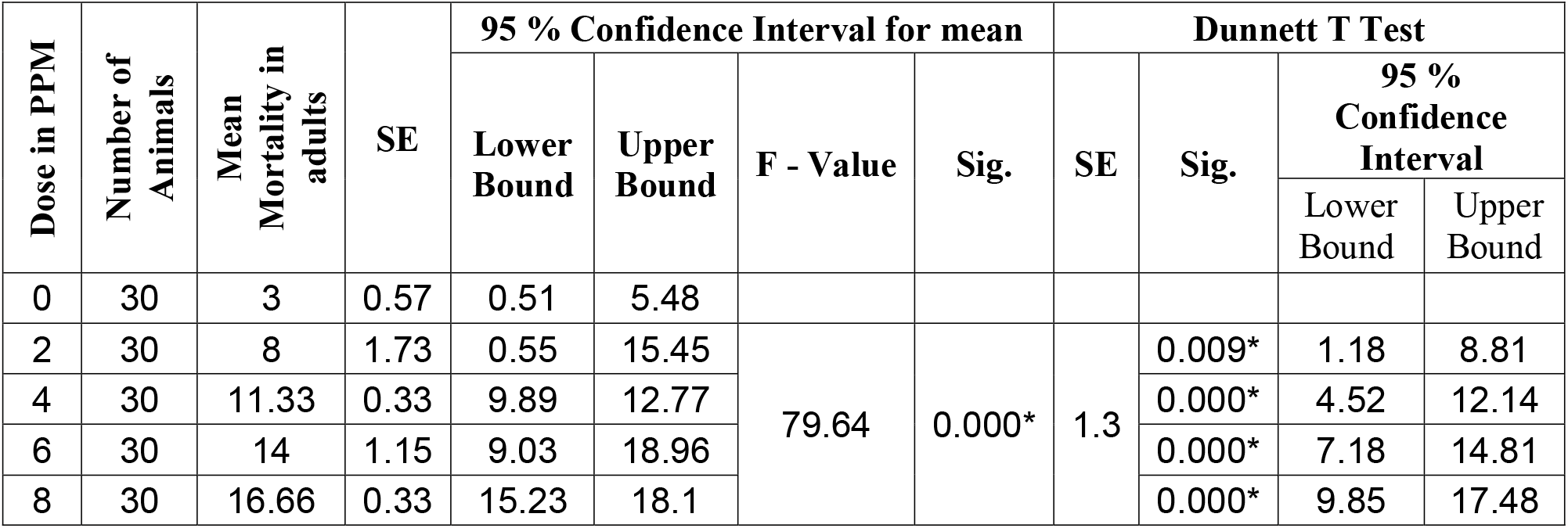
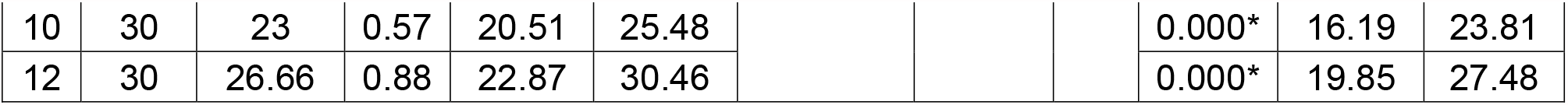
Results of One Way ANOVA and Dunnet T test for Acute toxicity in *Perna viriids*

The results of the regression determined the median lethal concentrations (LC_50_) as 6.10 PPM. Parameter estimates for analysis is presented in **Table 2**. Dose-response curve (R^2^ linear = 0.963) shows that, as the dose of the experimental groups increases, there is a corresponding increase in the mortality (**Fig. 1**).

**Table 2.** Parameter Estimates of Analysis (*Perna viridis*)

| Parameter Estimates |  |  |  |  |  |  |  |
| --- | --- | --- | --- | --- | --- | --- | --- |
|  | Parameter | Estimate | Std. Error | Z | Sig. | 95% Confidence Interval |  |
|  |  |  |  |  |  | Lower Bound | Upper Bound |
| <sup>a</sup> | Concentration | .214 | .028 | 7.733 | .000 | .160 | .269 |
|  | Intercept | -1.372 | .198 | -6.928 | .000 | -1.570 | -1.174 |
| a. model: (p) = Intercept + BX |  |  |  |  |  |  |  |

**Fig. 1.**
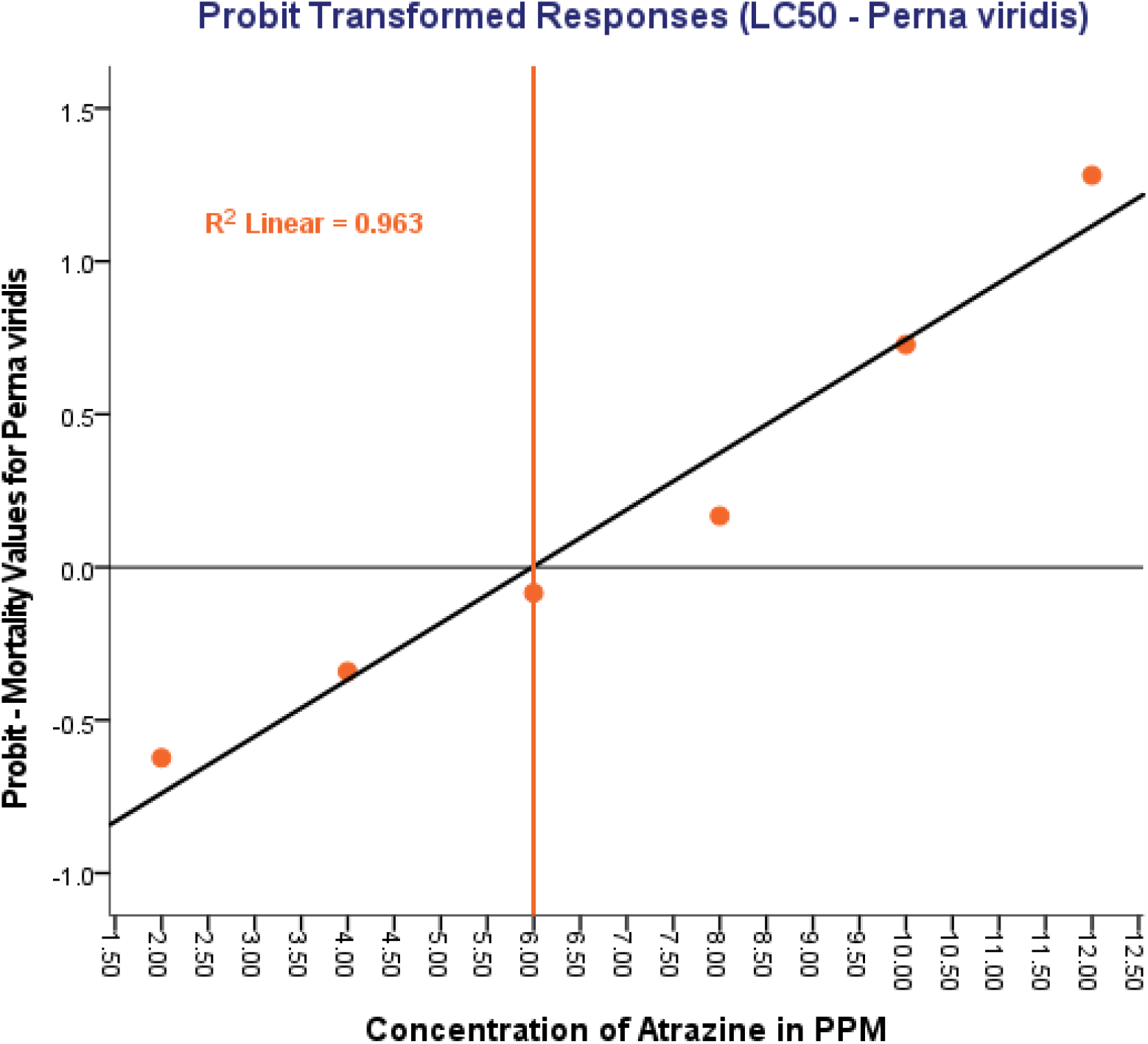
Median Lethal Concentration (LC_50_) determination by analysis in *Perna viridis*

#### Viable Hemocytes Count (VHC)

Acute toxicity experiments were carried out for over a period of 14 days to evaluate the percent viability and non-viability. Unstained (viable) and stained (nonviable) cells were identified by Tryphan Blue Exclusion Assay. The average count of viable hemocytes, nonviable hemocytes, total hemocytes and percentage of viability is presented in **Table 3**. The counting of unstained cells (viable hemocytes) was carried out on 2, 4, 6, 8, 10, 12 and 14 days for the control and experimental groups, percentage of the viability was calculated and presented in **Fig. 2**. The VHC and percentage viability in control group found to be between 94.34 (2.034 × 10^7^ ml^-1^) on 2^nd^ day to 92.34 (1.952 × 10^7^ ml^-1^) on 14^th^ day. No significant difference was detected within the control group. In experimental group of lowest observed effective level (LOEL) of 0.25 PPM the number of viable cells decreased gradually from 90.5 (1.96 × 10^7^ ml^-1^) on 2^nd^ day to 85.31 (1.85 × 10^7^ ml^-1^) on 14^th^ day. In second experimental group with 0.50 PPM of atrazine the percentage of viable cells decreased from 87.6 (1.92 × 10^7^ ml^-1^) to 84.47 (1.72 × 10^7^ ml^-1^) on day 2 and day14 respectively. The trend of decrease in percentage of viable hemocytes has continued with the corresponding increase in the doses of 0.75 and 1.00 PPM. Experimental group with highest dose treatment of 1.25 PPM, percentage of viable hemocytes was recorded to be 80.86 (1.68× 10^7^ ml^-1^) on 2^nd^ day and further decreased to reach a minimum of 74.51 (1.42 × 10^7^ ml^-1^) on 14^th^ day. The detailed data of percentage of viability is presented **in Fig.3**

**Table 3.** Viable (VHC), Non-viable (NVHC), Total hemocytes (THC) (cells ml^-1^) and percentage of viability in *Perna viridis*, (14 Days atrazine Exposure)

|  |  |  | Duration of the Exposure (In Days) |  |  |  |  |  |  |
| --- | --- | --- | --- | --- | --- | --- | --- | --- | --- |
|  |  |  | 2 | 4 | 6 | 8 | 10 | 12 | 14 |
| Atrazine Dose Exposure (in PPM) | 1.25 | VH C | $1.68 \times 10^7$ | $1.63 \times 10^7$ | $1.59 \times 10^7$ | $1.57 \times 10^7$ | $1.54 \times 10^7$ | $1.51 \times 10^7$ | $1.42 \times 10^7$ |
| | | NVH C | $0.40 \times 10^7$ | $0.40 \times 10^7$ | $0.40 \times 10^7$ | $0.40 \times 10^7$ | $0.42 \times 10^7$ | $0.44 \times 10^7$ | $0.49 \times 10^7$ |
| | | THC | $2.08 \times 10^7$ | $2.04 \times 10^7$ | $1.99 \times 10^7$ | $1.97 \times 10^7$ | $1.96 \times 10^7$ | $1.95 \times 10^7$ | $1.91 \times 10^7$ |
|  |  | % | **80.86** | **80.16** | **79.82** | **79.8** | **78.75** | **77.3** | **74.51** |
| | 1.00 | VHC | $1.81 \times 10^7$ | $1.78 \times 10^7$ | $1.73 \times 10^7$ | $1.67 \times 10^7$ | $1.65 \times 10^7$ | $1.60 \times 10^7$ | $1.58 \times 10^7$ |
| | | NVH C | $0.31 \times 10^7$ | $0.33 \times 10^7$ | $0.34 \times 10^7$ | $0.35 \times 10^7$ | $0.38 \times 10^7$ | $0.39 \times 10^7$ | $0.40 \times 10^7$ |
| | | THC | $2.12 \times 10^7$ | $2.10 \times 10^7$ | $2.07 \times 10^7$ | $2.02 \times 10^7$ | $2.03 \times 10^7$ | $2.00 \times 10^7$ | $2.00 \times 10^7$ |
|  |  | % | **85.55** | **84.41** | **83.4** | **82.51** | **81.1** | **80.27** | **79.76** |
| | 0.75 | VHC | $1.94 \times 10^7$ | $1.93 \times 10^7$ | $1.85 \times 10^7$ | $1.76 \times 10^7$ | $1.72 \times 10^7$ | $1.68 \times 10^7$ | $1.72 \times 10^7$ |
| | | NVH<br>C | $0.28 \times 10^7$ | $0.29 \times 10^7$ | $0.32 \times 10^7$ | $0.35 \times 10^7$ | $0.36 \times 10^7$ | $0.37 \times 10^7$ | $0.40 \times 10^7$ |
| | | THC | $2.22 \times 10^7$ | $2.22 \times 10^7$ | $2.18 \times 10^7$ | $2.10 \times 10^7$ | $2.08 \times 10^7$ | $2.06 \times 10^7$ | $2.12 \times 10^7$ |
|  |  | % | **86.76** | **87.48** | **85.2** | **83.41** | **82.66** | **81.82** | **81.1** |
| | 0.50 | VHC | $1.92 \times 10^7$ | $1.94 \times 10^7$ | $1.89 \times 10^7$ | $1.84 \times 10^7$ | $1.79 \times 10^7$ | $1.77 \times 10^7$ | $1.77 \times 10^7$ |
| | | NVH<br>C | $0.27 \times 10^7$ | $0.23 \times 10^7$ | $0.28 \times 10^7$ | $0.30 \times 10^7$ | $0.32 \times 10^7$ | $0.33 \times 10^7$ | $0.33 \times 10^7$ |
| | | THC | $2.19 \times 10^7$ | $2.17 \times 10^7$ | $2.16 \times 10^7$ | $2.14 \times 10^7$ | $2.11 \times 10^7$ | $2.10 \times 10^7$ | $2.10 \times 10^7$ |
|  |  | % | **87.6** | **89.39** | **87.22** | **86.17** | **85.01** | **84.35** | **84.47** |
| | 0.25 | VHC | $1.96 \times 10^7$ | $2.05 \times 10^7$ | $2.02 \times 10^7$ | $1.99 \times 10^7$ | $1.94 \times 10^7$ | $1.91 \times 10^7$ | $1.85 \times 10^7$ |
| | | NVH<br>C | $0.20 \times 10^7$ | $0.19 \times 10^7$ | $0.20 \times 10^7$ | $0.22 \times 10^7$ | $0.24 \times 10^7$ | $0.28 \times 10^7$ | $0.32 \times 10^7$ |
| | | THC | $2.17 \times 10^7$ | $2.24 \times 10^7$ | $2.22 \times 10^7$ | $2.22 \times 10^7$ | $2.19 \times 10^7$ | $2.20 \times 10^7$ | $2.17 \times 10^7$ |
|  |  | % | **90.5** | **91.53** | **90.89** | **89.9** | **89.13** | **87.07** | **85.31** |
| | Control | VHC | $2.03 \times 10^7$ | $1.98 \times 10^7$ | $1.98 \times 10^7$ | $1.96 \times 10^7$ | $1.94 \times 10^7$ | $1.96 \times 10^7$ | $1.95 \times 10^7$ |
| | | NVH<br>C | $0.12 \times 10^7$ | $0.13 \times 10^7$ | $0.14 \times 10^7$ | $0.13 \times 10^7$ | $0.14 \times 10^7$ | $0.16 \times 10^7$ | $0.16 \times 10^7$ |
| | | THC | $2.16 \times 10^7$ | $2.11 \times 10^7$ | $2.12 \times 10^7$ | $2.09 \times 10^7$ | $2.08 \times 10^7$ | $2.12 \times 10^7$ | $2.11 \times 10^7$ |
|  |  | % | **94.34** | **93.66** | **93.21** | **93.69** | **93.07** | **92.47** | **92.34** |

**Fig. 2.**
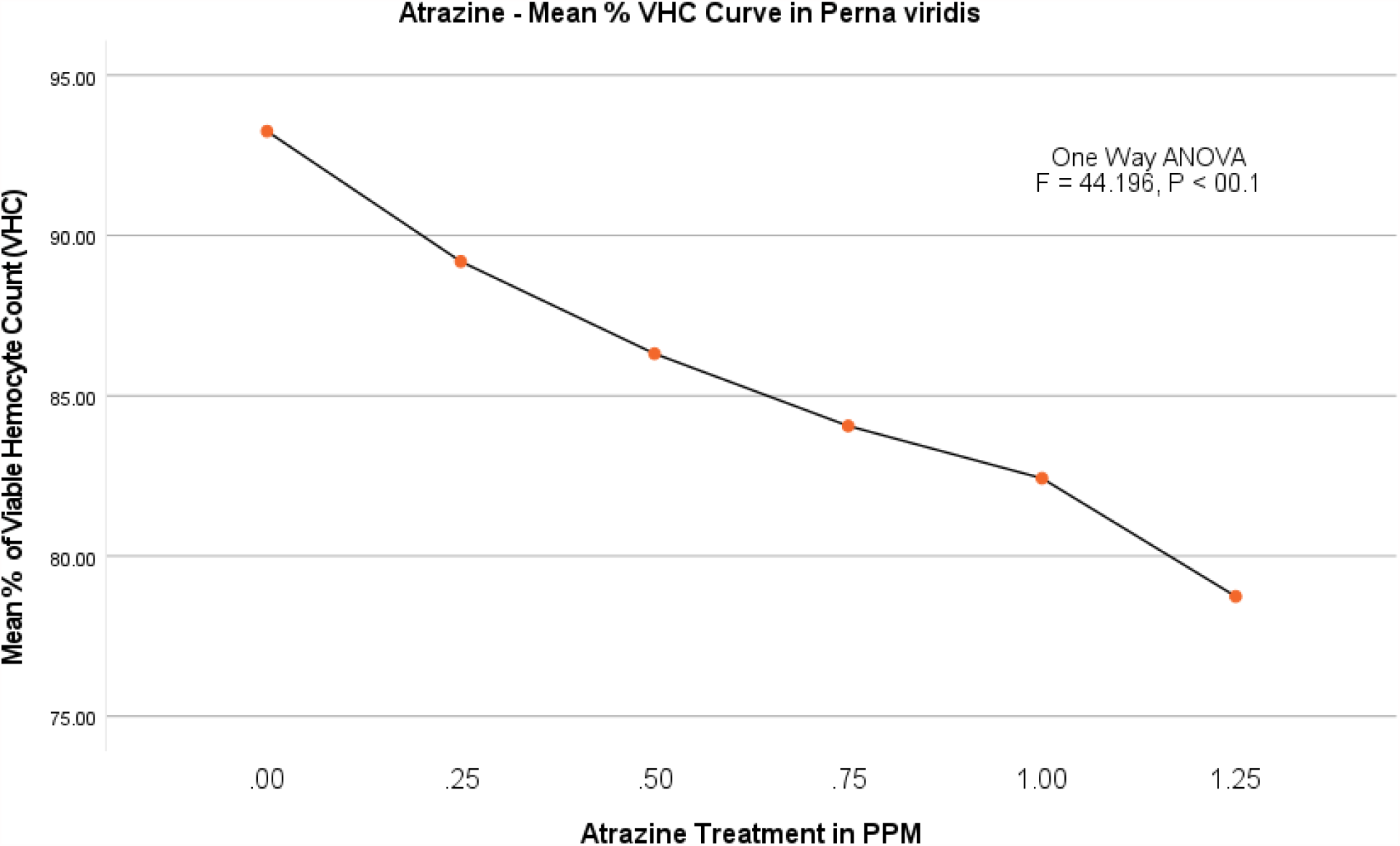
Atrazine dose – VHC response curve in *Perna viridis*

**Fig. 3.**
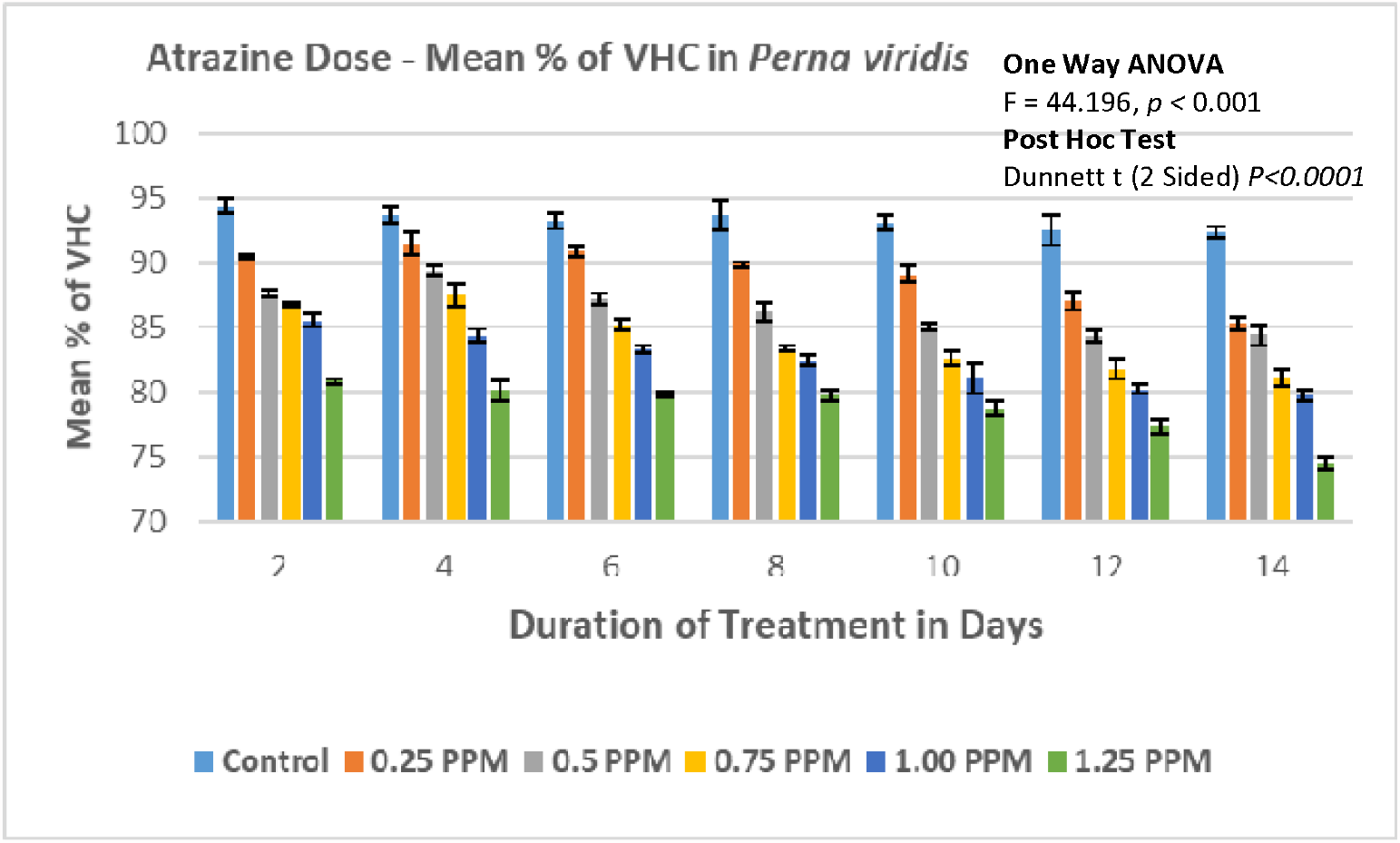
Percentage of viability of hemocytes in *Perna viridis*.

One Way ANOVA test was used to evaluate the level of significance between the viable hemocytes Count in the Control group and the experimental groups. This test shows the significant difference in viability between the groups (F = 28.566, P < 0.001). Further, multiple comparisons performed by Dunnett T test, the mean difference is significant (P < 0.0003) between the groups with 95 % confidence interval.

In order to rule out the errors in the assay technique, we have also evaluated the count and percentage of the non-viable hemocytes. There was a drastic increase in the non–viable hemocytes in *Perna viridis*. However, nonviable hemocytes count is inevitable to calculate total hemocytes. The dark blue stained nonviable hemocytes were recorded and their percentage calculated. In control group percentage of nonviable hemocytes on the 2^nd^ day of the experiment were calculated 5.781 (0.122 × 10^7^ ml^-1^) and on the last day of the experiment i.e. 14th day 7.66 (0.160 × 10^6^). There is a just marginal difference over a period of 14 days. However, in experimental groups steady increase in the nonviable hemocytes is evident (**Fig.4**). In case of lowest observed effective level (LOEL) of 0.25 PPM on 2^nd^ day percentage of nonviable hemocytes was 9.18 (0.206 × 10^6^) and on 14^th^ day percentage gone up to 14.68 (0.318 × 10^6^). Correspondingly, there is an incremental increase in percent nonviability in experimental groups of 0.50, 0.75, 1.00 PPM. Finally at experimental group with 1.25 PPM challenge, the percentage of non-viability recorded as 18.91 (0.398 × 10^7^ ml^-1^) on 2^nd^ day itself and 24.47 (0.486× 10^7^ ml^-1^) One Way ANOVA test was repeated by using IBM SPSS 24, which showed significant difference between the groups (F = 47.118, P < 0.001) in nonviable hemocytes. The percentage of nonviable cells is presented in **Fig.5**. Further, the multiple comparision by using Dunnett T test, the mean difference is significant for all the experimental groups when individually compared with control group (P < 0.001) with 95 % confidence interval.

**Fig. 4.**
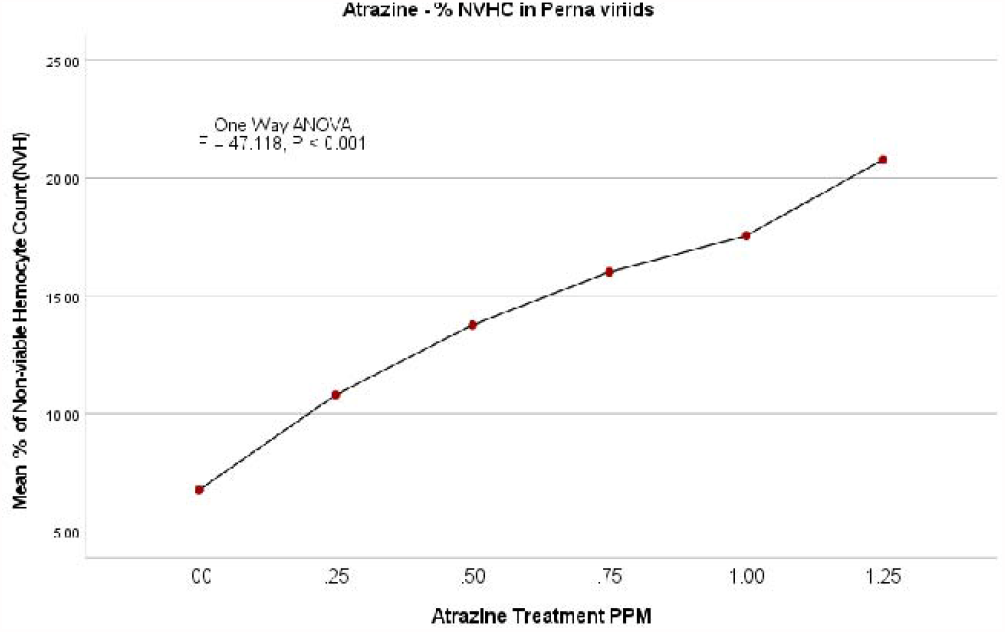
Atrazine dose – NVHC response curve in *Perna viridis*

**Fig. 5.**
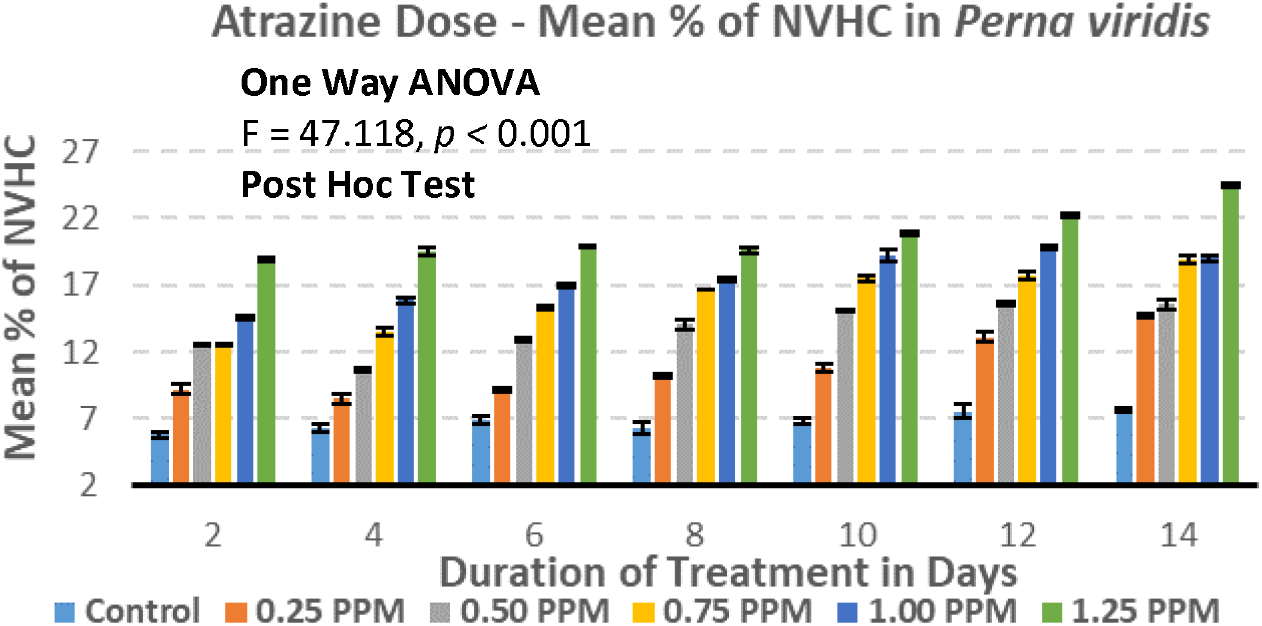
Percentage of nonviability of hemocytes in *Perna viridis*

### Paphia malabarica

#### Acute toxicity studies

Even in *Paphia malabarica*, until now no acute toxicological studies have been carried out and reported. In this article we are reporting the median lethal concentration (LC_50_) of atrazine and lowest observed effective level (LOAL) were evaluated by conducting 96 hour LC_50_ experiments in triplicates as described for the *Perna viridis*. Mortality recorded for every 24, 48, 72 and 96 hours. Results of one way ANOVA showed significant difference in mortality between control and experimental groups with F = 138.6 and P < 0.0001, Dunnett T test results of multiple comparisons also show significant difference for all experimental groups when compared with control group (P < 0.0001). The LOEL in this species also found to be 2.00 PPM (P < 0.004) with 95 % confidence intervals (2.24, 11.76). The results are presented in **Table 4**.

**Table 4.** Results of One Way ANOVA and Dunnet T test for Acute toxicity in *Paphia malabarica*

| Dose in PPM | Number of Animals | Mean Mortality in adults | SE | 95 % Confidence Interval for mean |  |  |  | Dunnett T Test |  |  |  |
| --- | --- | --- | --- | --- | --- | --- | --- | --- | --- | --- | --- |
|  |  |  |  | Lower Bound | Upper Bound | F - Value | Sig. | SE | Sig. | 95 % Confidence Interval |  |
|  |  |  |  |  |  |  |  |  |  | Lower Bound | Upper Bound |
| 0 | 30 | 8.66 | 0.33 | 7.23 | 10.1 |  |  |  |  |  |  |
| 2 | 30 | 15.66 | 0.66 | 12.8 | 18.53 | 138.6 | 0.000* | 1.6 | 0.004* | 2.24 | 11.76 |
| 4 | 30 | 21.66 | 0.33 | 20.23 | 23.1 |  |  |  | 0.000* | 8.24 | 17.76 |
| 6 | 30 | 27 | 0.57 | 25.51 | 30.48 |  |  |  | 0.000* | 14.58 | 24.08 |
| 8 | 30 | 34.66 | 2.6 | 23.46 | 45.87 |  |  |  | 0.000* | 21.24 | 30.76 |
| 10 | 30 | 41.66 | 0.33 | 40.23 | 43.1 |  |  |  | 0.000* | 28.24 | 37.76 |
| 12 | 30 | 45.66 | 1.2 | 40.49 | 50.83 |  |  |  | 0.000* | 32.24 | 41.76 |

The Regression test determined median lethal concentrations LC_50_ as 4.90 PPM. Parameter estimates for analysis and Dose-response curve (R^2^ linear = 0.996) are presented in **Table 5 & Fig.6**.

**Table 5.** Parameter Estimates of Analysis (*Paphia malabarica*)

| Parameter Estimates |  |  |  |  |  |  |  |
| --- | --- | --- | --- | --- | --- | --- | --- |
|  | Parameter | Estimate | Std. Error | Z | Sig. | 95% Confidence Interval |  |
|  |  |  |  |  |  | Lower Bound | Upper Bound |
| a | Concentration | .180 | .024 | 7.337 | .000 | .132 | .228 |
|  | Intercept | -.904 | .177 | -5.095 | .000 | -1.081 | -.726 |
| a. model: (p) = Intercept + BX |  |  |  |  |  |  |  |

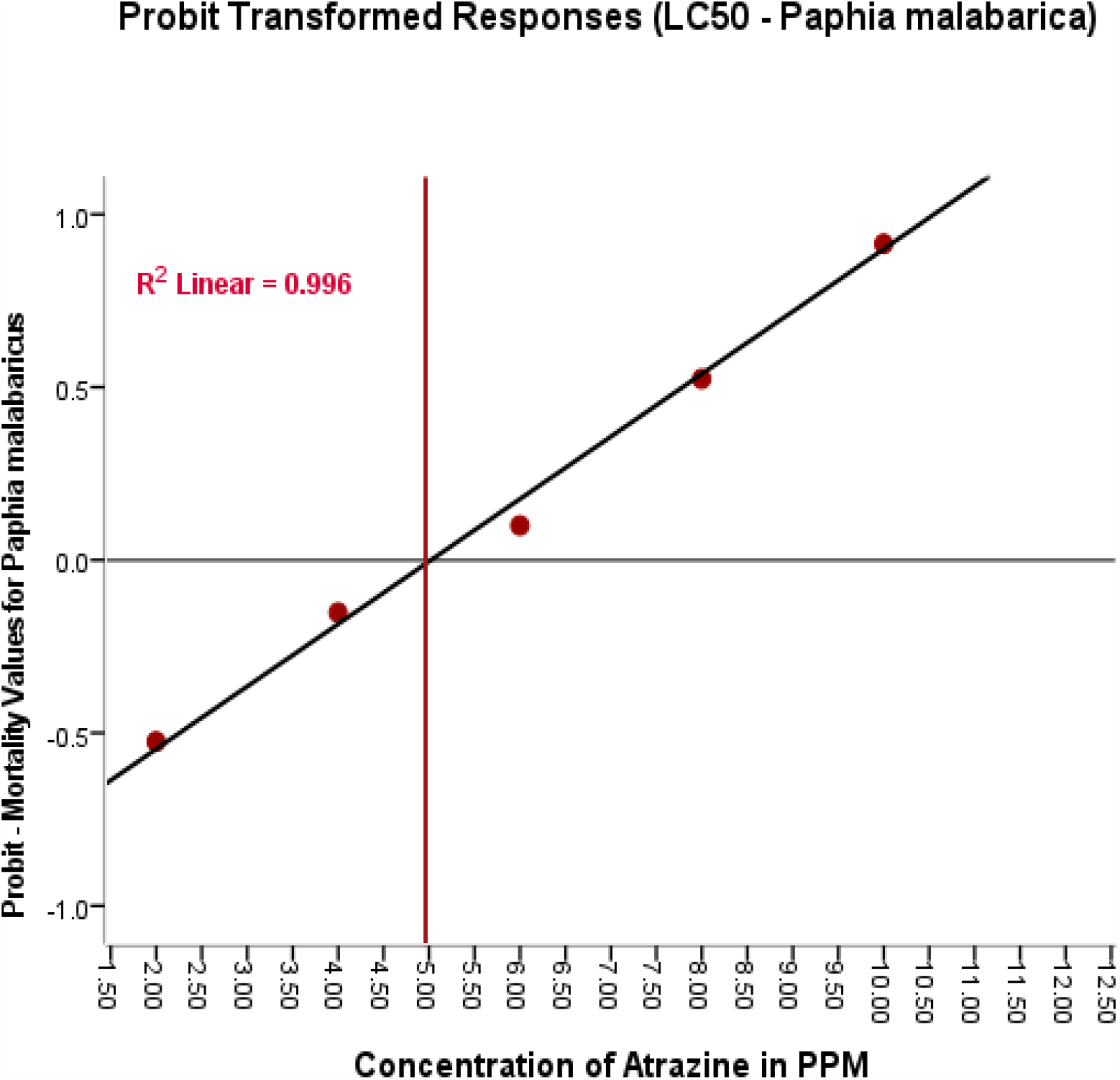

#### Viable Hemocytes Count (VHC)

The average count of viable hemocytes, nonviable hemocytes, total hemocytes and percentage of viability is presented in **Table 6**. The percentage of the viability was calculated and presented in **Fig.7**.The percentage of viability in control group found to be between 95.29 (1.54 × 10^7^ ml^-1^) on 2^nd^ day to 94.19 (1.59 × 10^7^ ml^-1^) on 14^th^ day. As expected, no significant difference was found in control group over a period of 14 days. In experimental group of lowest observed effective level (LOEL) of 0.20 PPM the number of viable cells steadily decreased from 90.56 (1.52 × 10^7^ ml^-1^) on 2^nd^ day to 84.69 (1.36 × 10^7^ ml^-1^) on 14^th^ day. The trend of decrease in percentage of viable hemocytes has continued with the corresponding increase in the doses of 0.40, 0.60 and 0.80 PPM. Finally, experimental group with highest dose treatment of 1.00 PPM, percentage of viable hemocytes was recorded to be 85.69 (1.35 × 10^7^ ml^-1^) on 2^nd^ day, which reached to a minimum of 78.39 (1.23 × 10^7^ ml^-1^) on 14^th^ day **Fig.8**. Data shows significant decrease in the percentage of viability in treated and untreated clams.

**Table 6.** Viable (VHC), Non-viable (NVHC), Total hemocytes (THC) (cells ml^-1^) and percentage of viability in *Paphia malabarica* (14 Days atrazine Exposure)

|  |  |  | Duration of the Exposure (In Days) |  |  |  |  |  |  |
| --- | --- | --- | --- | --- | --- | --- | --- | --- | --- |
|  |  |  | 2 | 4 | 6 | 8 | 10 | 12 | 14 |
| Dose Exposure (In PPM) | 1 | VHC | $1.35 \times 10^7$ | $1.34 \times 10^7$ | $1.31 \times 10^7$ | $1.31 \times 10^7$ | $1.29 \times 10^7$ | $1.26 \times 10^7$ | $1.23 \times 10^7$ |
| | | NVHC | $0.23 \times 10^7$ | $0.25 \times 10^7$ | $0.27 \times 10^7$ | $0.29 \times 10^7$ | $0.31 \times 10^7$ | $0.31 \times 10^7$ | $0.34 \times 10^7$ |
| | | THC | $1.58 \times 10^7$ | $1.58 \times 10^7$ | $1.59 \times 10^7$ | $1.6 \times 10^7$ | $1.59 \times 10^7$ | $1.58 \times 10^7$ | $1.56 \times 10^7$ |
|  |  | % | **85.68** | **84.46** | **82.79** | **82.11** | **80.82** | **80.08** | **78.39** |
| | 0.8 | VHC | $1.31 \times 10^7$ | $1.32 \times 10^7$ | $1.31 \times 10^7$ | $1.30 \times 10^7$ | $1.30 \times 10^7$ | $1.31 \times 10^7$ | $1.29 \times 10^7$ |
| | | NVHC | $0.22 \times 10^7$ | $0.24 \times 10^7$ | $0.26 \times 10^7$ | $0.27 \times 10^7$ | $0.28 \times 10^7$ | $0.28 \times 10^7$ | $0.31 \times 10^7$ |
| | | THC | $1.54 \times 10^7$ | $1.56 \times 10^7$ | $1.56 \times 10^7$ | $1.57 \times 10^7$ | $1.59 \times 10^7$ | $1.59 \times 10^7$ | $1.59 \times 10^7$ |
|  |  | % | **85.43** | **84.86** | **83.50** | **82.89** | **82.23** | **82.26** | **80.67** |
| | 0.6 | VHC | $1.40 \times 10^7$ | $1.37 \times 10^7$ | $1.34 \times 10^7$ | $1.35 \times 10^7$ | $1.31 \times 10^7$ | $1.30 \times 10^7$ | $1.26 \times 10^7$ |
| | | NVHC | $0.19 \times 10^7$ | $0.20 \times 10^7$ | $0.23 \times 10^7$ | $0.26 \times 10^7$ | $0.27 \times 10^7$ | $0.29 \times 10^7$ | $0.30 \times 10^7$ |
| | | THC | $1.59 \times 10^7$ | $1.58 \times 10^7$ | $1.57 \times 10^7$ | $1.61 \times 10^7$ | $1.57 \times 10^7$ | $1.59 \times 10^7$ | $1.56 \times 10^7$ |
|  |  | % | **88.03** | **87.05** | **85.09** | **84.07** | **83.11** | **81.85** | **80.94** |
| | 0.4 | VHC | $1.43 \times 10^7$ | $1.41 \times 10^7$ | $1.36 \times 10^7$ | $1.35 \times 10^7$ | $1.34 \times 10^7$ | $1.33 \times 10^7$ | $1.33 \times 10^7$ |
| | | NVHC | $0.17 \times 10^7$ | $0.19 \times 10^7$ | $0.22 \times 10^7$ | $0.24 \times 10^7$ | $0.25 \times 10^7$ | $0.25 \times 10^7$ | $0.26 \times 10^7$ |
| | | THC | $1.60 \times 10^7$ | $1.60 \times 10^7$ | $1.58 \times 10^7$ | $1.59 \times 10^7$ | $1.54 \times 10^7$ | $1.58 \times 10^7$ | $1.58 \times 10^7$ |
|  |  | % | **89.49** | **87.97** | **85.94** | **84.78** | **87.27** | **84.07** | **83.71** |
| | 0.2 | VHC | $1.52 \times 10^7$ | $1.46 \times 10^7$ | $1.45 \times 10^7$ | $1.42 \times 10^7$ | $1.38 \times 10^7$ | $1.37 \times 10^7$ | $1.36 \times 10^7$ |
| | | NVHC | $0.16 \times 10^7$ | $0.17 \times 10^7$ | $0.19 \times 10^7$ | $0.19 \times 10^7$ | $0.21 \times 10^7$ | $0.23 \times 10^7$ | $0.23 \times 10^7$ |
| | | THC | $1.67 \times 10^7$ | $1.62 \times 10^7$ | $1.64 \times 10^7$ | $1.61 \times 10^7$ | $1.59 \times 10^7$ | $1.60 \times 10^7$ | $1.60 \times 10^7$ |
|  |  | % | **90.57** | **89.77** | **88.50** | **88.19** | **87.01** | **85.76** | **85.45** |
| | Control | VHC | $1.54 \times 10^7$ | $1.55 \times 10^7$ | $1.53 \times 10^7$ | $1.5 \times 10^7$ | $1.56 \times 10^7$ | $1.59 \times 10^7$ | $1.59 \times 10^7$ |
| | | NVHC | $0.08 \times 10^7$ | $0.08 \times 10^7$ | $0.09 \times 10^7$ | $0.10 \times 10^7$ | $0.11 \times 10^7$ | $0.10 \times 10^7$ | $0.10 \times 10^7$ |
| | | THC | $1.61 \times 10^7$ | $1.63 \times 10^7$ | $1.63 \times 10^7$ | $1.65 \times 10^7$ | $1.67 \times 10^7$ | $1.69 \times 10^7$ | $1.68 \times 10^7$ |
|  |  | % | **95.29** | **94.85** | **94.22** | **93.83** | **93.31** | **94.07** | **94.20** |

**Fig. 7.**
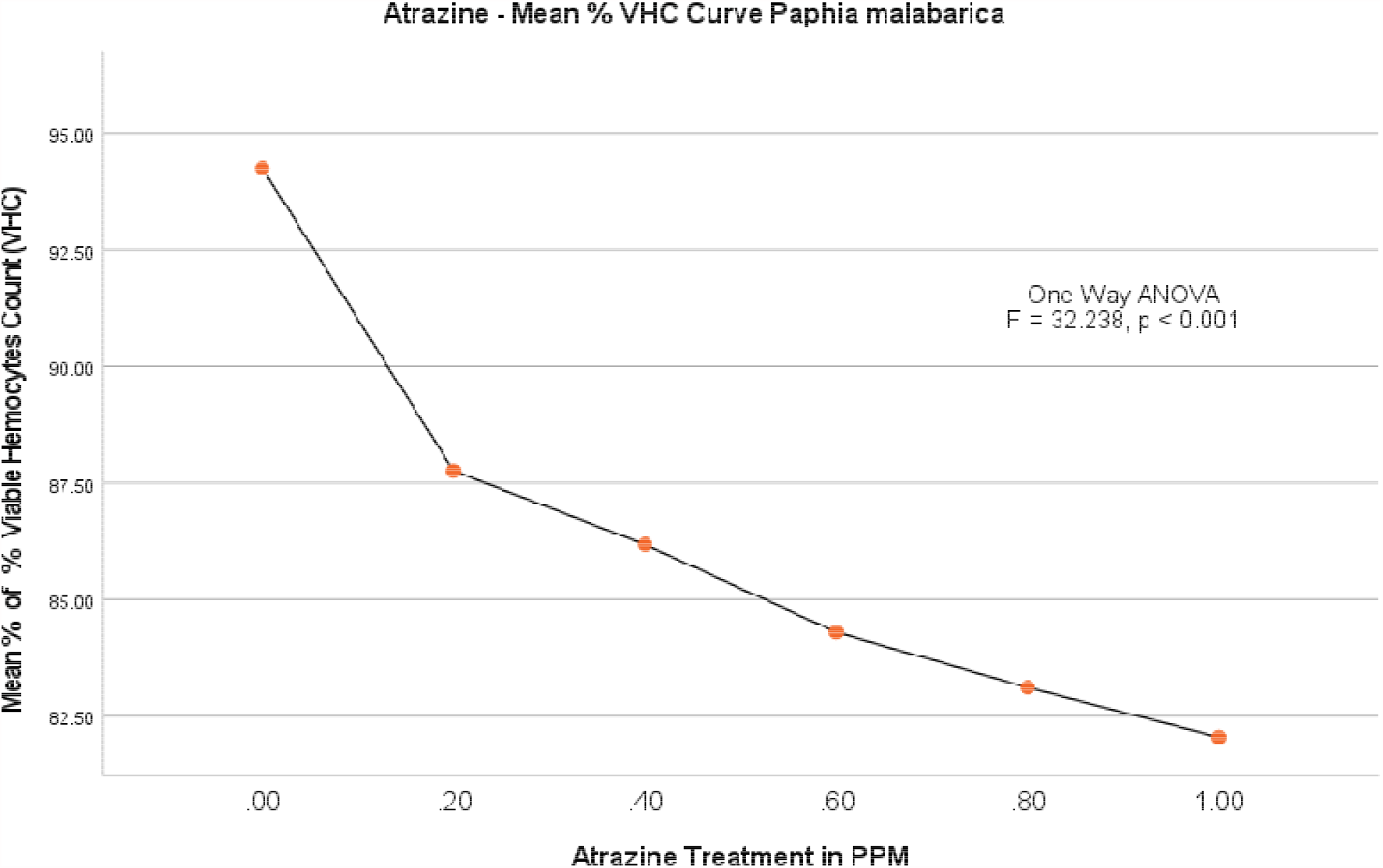
Atrazine dose – VHC response curve in *Paphia malabarica*

**Fig. 8.**
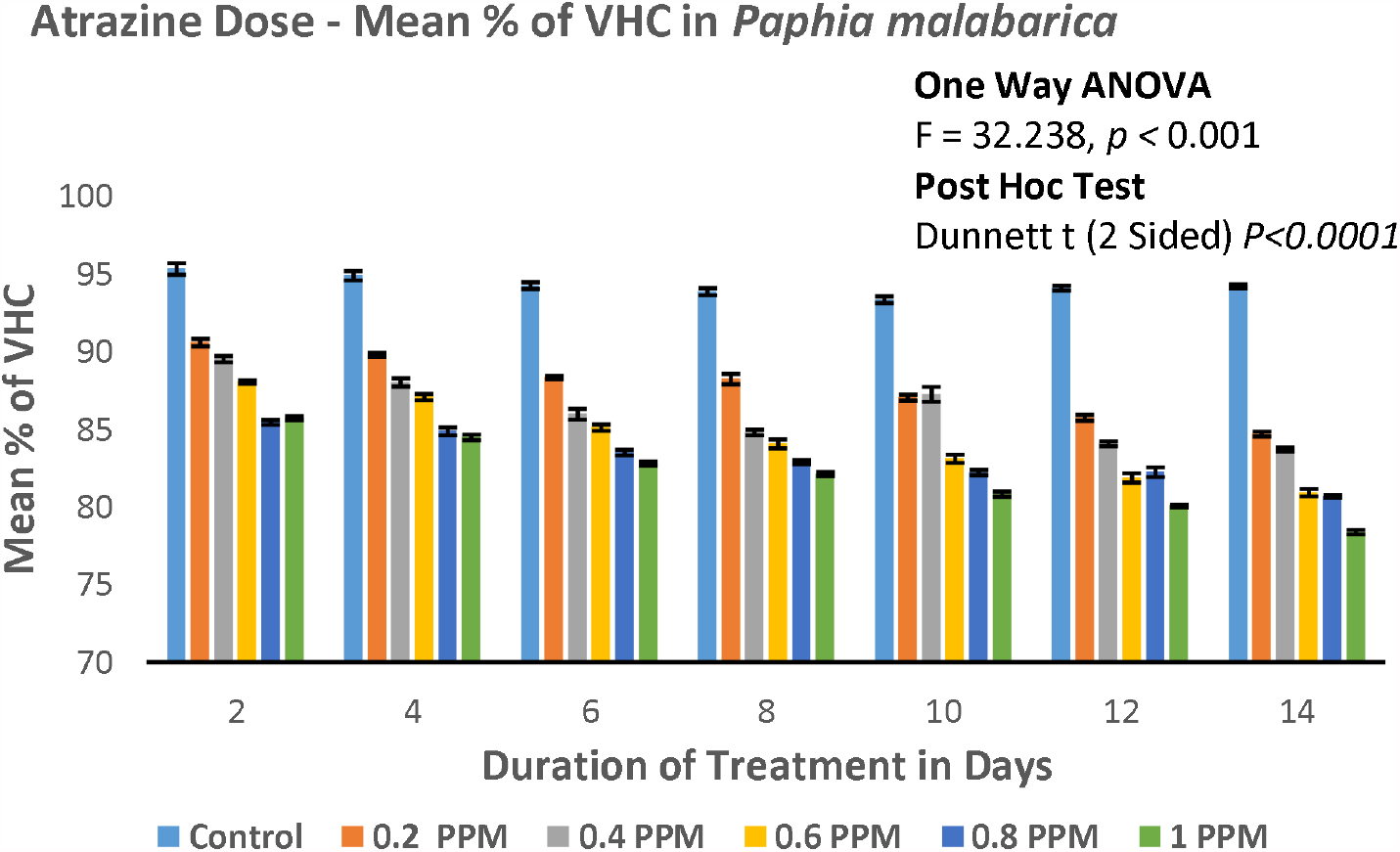
Percentage of viability of hemocytes in *Paphia malabarica*

One-Way ANOVA test showed significant difference in viability between the groups (F = 32.238, P < 0.001) but no significant difference within the groups. Dunnett T test, the mean difference is significant (P < 0.05) with 95 % confidence interval.

#### Non-viable hemocytes count (NVHC)

In control group percentage of nonviable hemocytes on the 2^nd^ day of the experiment were calculated 5.781 (0.122 × 10^7^ ml^-1^) and on the last day of the experiment i.e. 14th day 7.66 (0.160 × 10^6^). In control group, there is a just marginal difference over a period of 14 days. In case of lowest observed effective level (LOEL) of 0.20 PPM on 2^nd^ day percentage of non-viable hemocytes was 9.43 (0.16 × 10^7^ ml^-1^) and on 14^th^ day percentage gone up to 14.55 (0.23 × 10^7^ ml^-1^). Correspondingly, there is a significant progressive increase in percentage of non-viability in experimental groups of 0.40, 0.60, 0.80. Experimental group with highest treatment 1.00 PPM challenge, the percentage of non-viability recorded 14.30 (0.23 × 10^7^ ml^-1^) on 2^nd^ day itself and 21.6 (0.34 × 10^7^ ml^-1^). In case of non-viable hemocytes there is a significant progressive increase.

One Way ANOVA test was repeated by using IBM SPSS 24, which showed significant difference between the groups (F = 34.08, P < 0.001) in nonviable hemocytes. The results are presented in **Fig.9 and Fig. 10**. Further, the multiple comparisons by using Dunnett T test, the mean difference is significant at P < 0.001 with 95 % confidence interval.

**Fig. 9.**
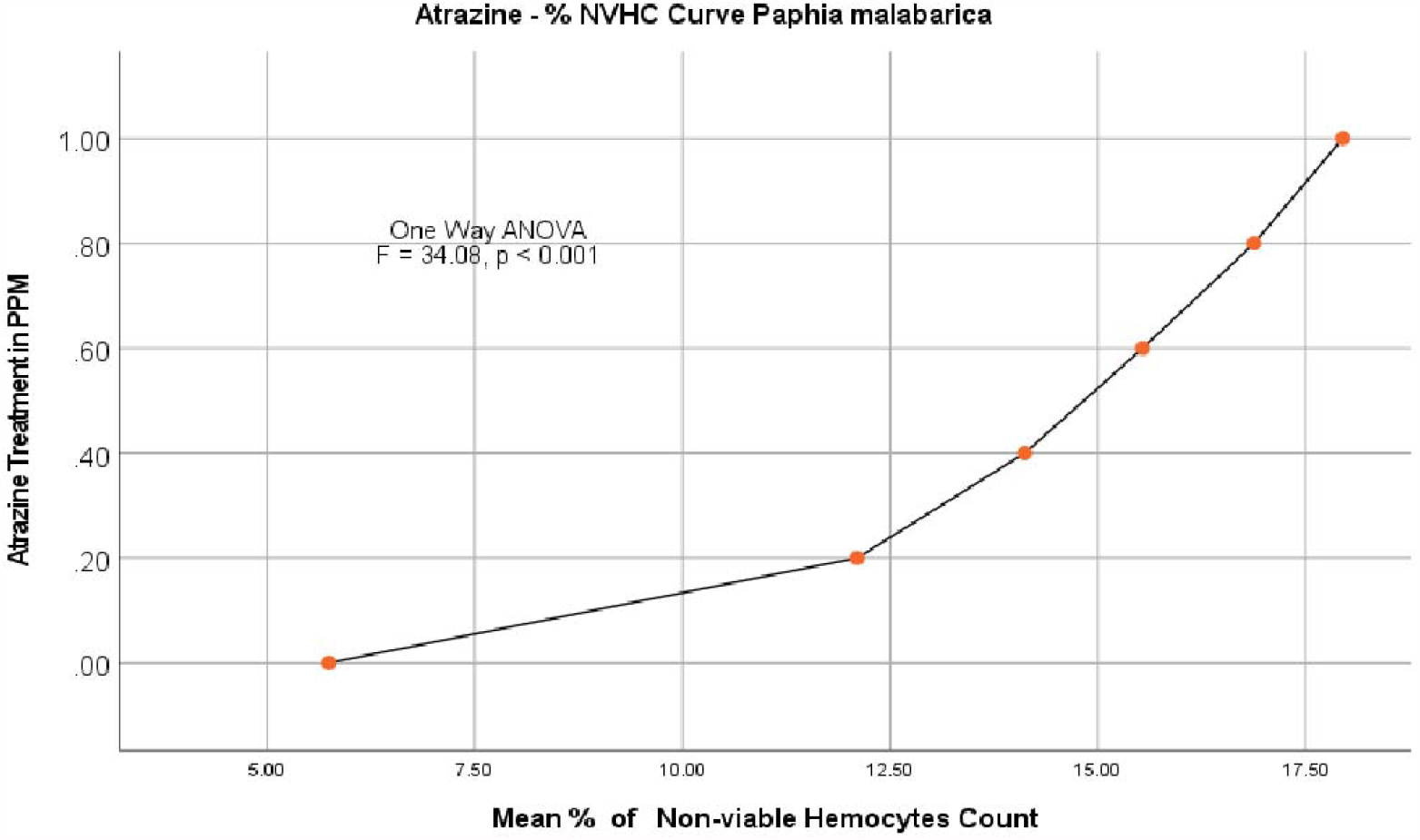
Atrazine dose – NVHC response curve in *Paphia malabarica*

**Fig. 10.**
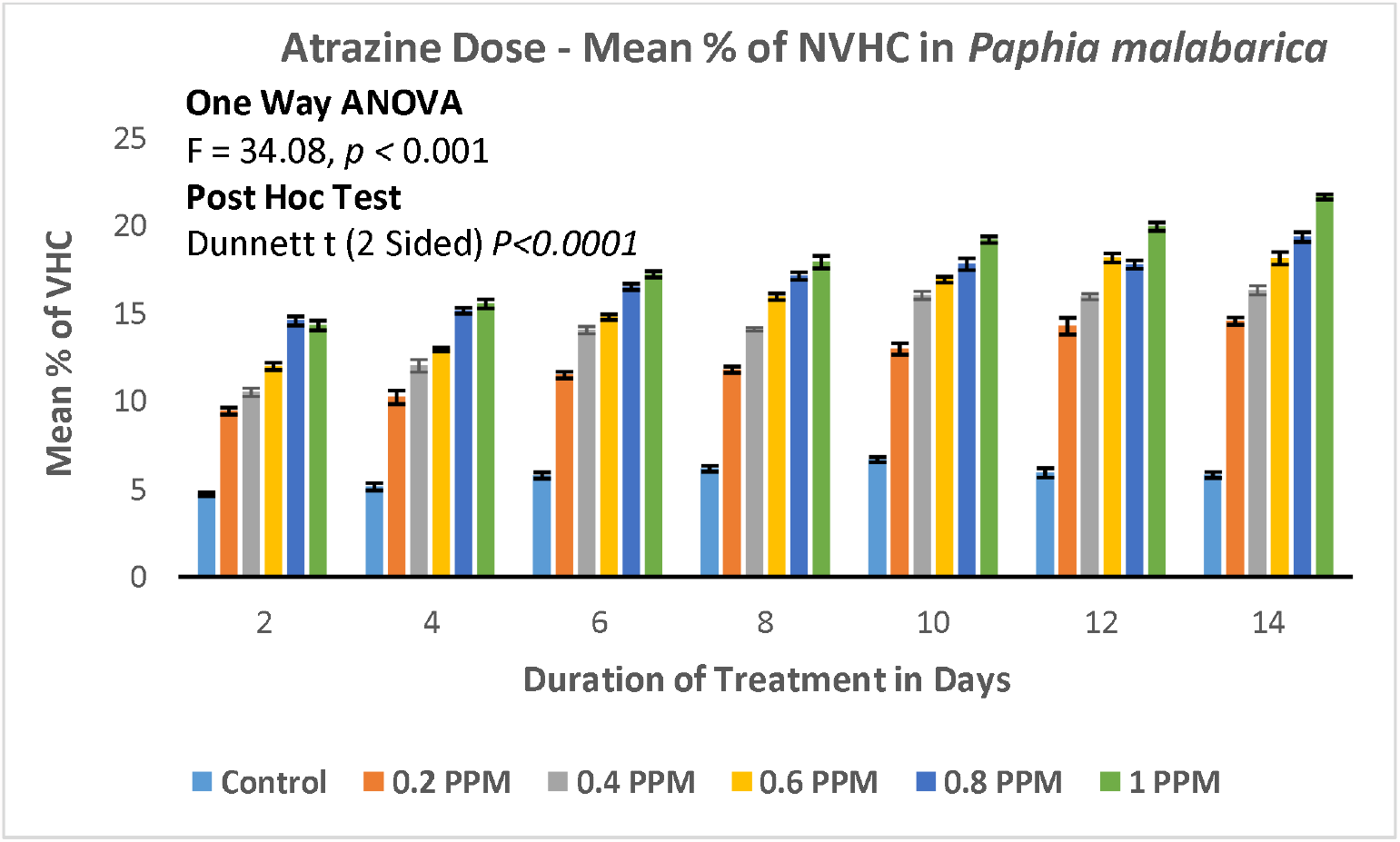
Percentage of Nonviability of hemocytes in *Paphia malabarica*

## Discussion

Environmental contaminants is one of the greatest challenges we are facing today, which is adversely affecting the quality of every ecosystem. Enormous number of chemicals are being synthesized on daily basis, most of them eventually released into environment without passing through any risk assessment and safety standards. Only few chemicals are subjected to proper risk assessment, others either get neglected or understudied. There is absolutely no match between the number of chemicals being synthesized and the studies on their risk assessment. Moreover, it is extremely difficult to subject millions of chemical to the toxicological studies. Nonetheless, few abundantly used chemicals such as pesticides and herbicides have received good scientific attention. Available scientific literature on the toxicity of chemicals suggests focused study on certain model organisms like mice, rat, rabbit and fishes while other organisms are mostly ignored. However, from the perspective of the method of application of some of the chemicals especially pesticides and herbicides, aquatic animals are more vulnerable. Long-term persistence of many of these chemicals in surface water pose greater challenges of acute and chronic toxicity to the aquatic animals. Amongst aquatic animals, bivalves are more vulnerable to the pollutants and environmental contaminants as their habitats receive maximum pollutant loads.

Toxicity of atrazine is reported from several studies carried out in fishes, amphibians and mammals. Majority of the In vivo and in vitro studies have reported deleterious effects of atrazine on the reproductive endocrinology and developmental toxicity whereas some studies have reported the toxicity related to cells of vertebrate immune system. Several manifestations of reproductive endocrinology such as reproductive organ anomalies, failure of implantations, and termination of early gestations, delayed onset of puberty in mice models were linked directly with the atrazine toxicity (Cummings et al., 2000; Stoker et. al. 2002; Laws et.al. 2003). There are few reports on the Immunotoxicity of atrazine on mice and other mammalian models wherein the atrazine found to significantly decrease the hematopoietic progenitor cells, reticulocytes and CD4^+^ lymphocytes (Pistl et al., 2002; Whale et al., 2003; Filipov et al., 2005). In addition, atrazine also reported to have immunomodulatory functions by significantly increasing the natural killer cells, T cell proliferation, IgM secreting plasma cells, macrophage mediated phagocytic and cytolytic functions, thus affecting both innate and adaptive immunity (Whale et al., 2003; Rowe et.al. 2006).

In fishes, atrazine toxicity is associated secondary lymphoid organs in salmonid fishes like Coho salmon (*Oncorhynchus kisut*ch) and lake trout (*Salvelinus namaycush)* (Zeeman and Brindley, 1981). Some reports even suggests atrazine induced decrease in the number of circulating lymphocytes resulting in leucopenia and degeneration of macrophages in fishes (Zeeman and Brindley, 1981). Apart from immunotoxicity, the atrazine is also reported to have endocrine toxicity in catfishes (Opute et. al. 2021). In amphibians, the atrazine is reported to decrease the number of spleenocytes and leucocytes in *Rana pipines, Xenopus laevis* and *Rana sylvatica* (Christin et al. 2013; Kiesecker; 2002). Decrease in number of these cells seriously undermines innate and adaptive immunity in amphibians.

Apart from aforementioned studies, no other scientific literature on the immunotoxic potential of atrazine is available either on vertebrates or on invertebrates. However, immunotoxicity of other environmental contaminants such as pesticides, heavy metals, PCBs have been discussed elaborately in few invertebrates including bivalves (Renault, 2015; Ellis et al. 2011; Destoumieux-Garzón et al. 2020). Even molecular mechanisms involved also been discussed in few species (Gerdol, and Venier, 2015; Bachère et al 2015; Burgos-Aceves and Faggio, 2017; Teresa et al. 2021).

In the present study, we have evaluated the acute toxicity of atrazine on Perna viridis (Order Mytilida) and Paphia malabarica (Venerida), LC50 was also determined and we have also examined the effect of sub-lethal doses of atrazine on the viability of hemocytes. The results of the present study clearly indicate that the atrazine significantly decreases the number of hemocytes in bivalves.

The present study reports the immunotoxic effects of atrazine on the bivalves for the first time. This study is done by using Tryphan blue exclusion assay, which is simple, economical and reliable assay to evaluate the viability of hemocytes (Piccinini et al. 2017). This assay can be accomplished in a very short period of time and very convenient for the assessment of the large number of samples in a given period. In order to overcome the limitations associated with this technique, we have examined both viable and nonviable cells. Analysis by one way ANOVA showed significant decrease in the viable cells in experimental groups as compared to control group (P < 0.0001) and similarly, there is a proportionate increase in the percentage of nonviable cells in experimental groups as compared to control group (P < 0.0001) for both the species studied. Further, we have also examined the data by multiple comparisons by Dunnett T test to compare control group individually with each of the experimental groups, this test also found to be statistically significant (p < 0.001) for each experimental group. In *Perna viridis*, in control group, the percentage of viability calculated to be 94.34 on 2^nd^ day to 92.34 on 14^th^ day whereas in case of experimental group with highest concentration (1.25 PPM) the percent viability recorded as 80.86 on 2^nd^ day, which drastically decreased to 74.51 by 14^th^ day of atrazine exposure. The percent nonvaibility, on the other hand complemented to the percent viability, in control 5.78 (2^nd^ day) to 7.66 (14^th^ day) but in experimental group with highest concentration of 1.25 PPM the percent nonviability increased to 18.91 (2^nd^ day) to 24.47 (14^th^ day). In case of *Paphia malabarica*, similar results have been recorded, percent viability under control group found to be 95.29 (2^nd^ day) to 94.19 (14^th^ day) but decreased considerably to 85.69 (2^nd^ day) to 78.39 (14^th^ day). Concomitantly, percent nonviability in control group was 5.78 (2^nd^ day) to 7.66 (14^th^ day) but the percent of nonviability increased to 14.30 (2^nd^ day) to 21.6 (14^th^ day).

These results clearly indicate immuno-toxic potential of atrazine by decreasing the number of viable hemocytes.

## Conclusion

Present study is the first of its kind to investigate the immunotoxic potential of atrazine. The study used simple yet very reliable technique to examine the immunotoxic potential of atrazine. This study after detailed investigation suggests the immunotpxicity of atrazine by decreasing the viability of hemocytes, which are the principle cells of innate immunity in bivalves. Study done on two species belong to different two orders are consistent with the result.

## Acknowledgement

We are grateful to University Grants Commission (UGC), New Delhi for providing financial assistance vide order no. **MRP(S)-757/10-11/KAKA088/UGC-SWRO** to carry out this study. We are also indebted to Department of Collegiate Education, Bangalore for providing necessary laboratory facilities to complete this work.

## Declaration

I hereby declare that the authorship is given as per their contribution to the work. Dr Muhammed Zafar Iqbal conceived the idea and carried out experimental work. M.Azhar Iqbal Navalgund done statistical analysis of the data and helped in interpretation of data.

